# Expression of *Chlamydomonas reinhardtii* chloroplast diacylglycerol acyltransferase-3 is activated by light in concert with triacylglycerol accumulation

**DOI:** 10.1101/2021.01.17.426997

**Authors:** María de las Mercedes Carro, Débora Soto, Leandro Mamone, Gabriela Gonorazky, Carolina Bagnato, María Verónica Beligni

## Abstract

Considerable progress has been made towards the understanding of triacylglycerol (TAG) accumulation in algae. One key aspect is finding conditions that trigger TAG production without reducing cell division. Previously, we identified a soluble diacylglycerol acyltransferase (DGAT), related to plant DGAT3, with heterologous DGAT activity. In this work, we demonstrate that *Chlamydomonas reinhardtii* DGAT3 localizes to the chloroplast and its expression is activated by light, in correspondence with TAG accumulation. *Dgat3* mRNAs and TAGs increased in both wild type and starch-deficient cells grown with acetate upon transferring them from dark or low light to higher light. Light-activated DGAT3 expression and TAG accumulation depended on the preexisting levels of TAGs, suggesting the existence of a regulatory loop. These results indicate that DGAT3 could be responsible, at least in part, for light-dependent TAG accumulation. Moreover, our results present DGAT3 as a promising target of future studies oriented to the industrial applications of TAGs.

## Introduction

Current interest in the development of sustainable sources of liquid fuels and other industrial compounds has produced a remarkable expansion in the study of eukaryotic microalgae as feedstock for triacylglycerols (TAGs) (Banerjee et al., 2016; Li-Beisson et al., 2019; Sato et al., 2017; Xu et al., 2016). One of the most remarkable contributions to this field was the finding that neutral lipid generation in algae could be induced by stress and nutrient depletion (Allen et al., 2018; Pal et al., 2011; Rearte et al., 2018; Sharma et al., 2012; Yang et al., 2015). The best characterized condition is nitrogen deprivation, which stimulates TAG production in the majority of the algal species studied, independently of their taxonomic classification and lifestyles (Blaby et al., 2013; Martin et al., 2014; Msanne et al., 2012; Weng et al., 2014). Likewise, most of the enzymes that participate in TAG synthesis investigated to date are those that respond to a lack of nitrogen (Banerjee et al., 2017; Boyle et al., 2012; Chen et al., 2015; Guo et al., 2017; Xu et al., 2018). These enzymes are mostly homologs of proteins characterized in animals, yeast and plants that are involved in the conventional pathway carried out in the endoplasmic reticulum (ER)/cytosol (Bhatt-Wessel et al., 2018; Liu et al., 2011; Lung and Weselake, 2006; Shockey et al., 2006). Despite these advances, the drawback of any production scheme based on nitrogen deprivation is that TAG accumulation occurs in parallel to a decrease in algal cell division, which reduces total lipid productivity (Takeuchi and Benning, 2019). Hence, finding conditions that yield large amounts of TAGs during optimal growth is one of the big conundrums of this research area.

With this challenge in mind, we set out to search for novel enzymes that could participate in TAG synthesis in algae of diverse evolutionary origins. Using a HMMER iterative approach (Bagnato et al., 2017a), we identified a soluble diacylglycerol acyltransferase (DGAT) exclusive to green algae and moderately related to plant DGAT3 (Bagnato et al., 2017b). DGATs catalyze the formation of TAG by performing an acylation of a fatty acid (FA), most frequently activated as an acyl-CoA, onto the sn-3 position of a molecule of diacylglycerol (DAG) (Yen et al., 2008). Phylogenetic analysis showed that the DGAT3 clade shares a most recent ancestor with a group of uncharacterized proteins from cyanobacteria, suggesting a prokaryotic origin, whereas the canonical DGATs, DGAT1 and DGAT2, are only related to proteins from eukaryotes. In addition, most of the algal sequences within the DGAT3 group have predicted chloroplast localization (Bagnato et al., 2017b). In plants, DGAT3 has been characterized in only a few species, including *Arabidopsis thaliana, Vernicia fordii* (tung tree) and *Arachis hypogaea* (peanut) (Aymé et al., 2018; Cao et al., 2013; Chi et al., 2014; Hernández et al., 2012; Saha et al., 2006). DGAT3 protein sequences show very distinctive features, particularly in comparison with DGAT1 and DGAT2 (Cao, 2011; Liu et al., 2012; Turchetto-Zolet et al., 2011). The most conspicuous difference between DGAT3 and its counterparts is the fact that its sequence exhibits a carboxy-terminal iron-sulfur (2Fe-2S) cluster-binding domain that has a fold similar to that present in thioredoxins (Aymé et al., 2018; Bagnato et al., 2017b). Although the role of this domain in DGAT3 remains unknown, the majority of the homologs that have been studied to date act as low-potential electron carriers (Fukuyama, 2004; Zu et al., 2002).

The acylation activities of DGAT1 and DGAT2 have been widely described and reviewed (Yen et al., 2008). The conserved amino acids of these two protein families have been extensively studied in animals, yeast and plants and some of the residues that are important for catalysis have been determined in mutagenesis experiments (Liu et al., 2011; Lopes et al., 2014; Stone et al., 2006). In contrast, DGAT3 activity has only been demonstrated *in vitro* for the *A. thaliana*, *A. hypogaea* and *Chlamydomonas reinhardtii* proteins (Aymé et al., 2018; Bagnato et al., 2017b; Chi et al., 2014; Hernández et al., 2012; Saha et al., 2006) and little is known about the identity or location of the acylation domains. We previously reported that *C. reinhardtii* DGAT3 expressed in *Escherichia coli* produced and increase in TAGs in the presence of oleate (Bagnato et al., 2017b). In DGAT3 protein sequence, we identified three histidine-containing motifs similar to those that perform acylation in glycerol-3-phosphate acyltransferase (GPAT) and acylglycerol-3-phosphate acyltransferase (AGPAT) (Takeuchi and Reue, 2009; Yamashita et al., 2014), but those motifs have not been functionally analyzed.

The content of hydrophobic regions in DGAT3 is considerably lower than those of DGAT1 and DGAT2. Additionally, many of the hydrophobic regions of the latter two proteins translate into transmembrane segments, whereas DGAT3 appears to have no transmembrane domains. Most remarkably, in DGAT1 and DGAT2, the catalytic motifs either flank or are partially embedded in hydrophobic regions, whereas the putative catalytic motifs of *C. reinhardtii* DGAT3 are flanked by hydrophilic regions. All these observations led us to hypothesize that DGAT3 participates in a soluble TAG synthesis pathway in the chloroplast, where it could act as an electron carrier in addition to performing an acylation reaction (Bagnato et al., 2017b).

In this work, we demonstrate empirically that *C. reinhardii* DGAT3 is localized to the chloroplast and advance into its metabolic regulation. We demonstrate that DGAT3 expression is induced by light in the presence of acetate and that this induction occurs in correspondence with light-dependent TAG accumulation. Both DGAT3 expression and TAG levels increase upon shifting cells from dark or low light to higher light, albeit affected by the particularities of each strain. Altogether, our results suggest that light-dependent TAG accumulation could be carried out, at least in part, by the action of DGAT3. In addition to providing insights into the possible enzymatic components of light-activated TAG production, our results further advance into the relationship between light and carbon in regards to TAG metabolism.

## Results

Figure 1A shows a schematic representation of *C. reinhardtii* DGAT3 full-length protein. This polypeptide has a transit peptide of approximately 70 amino acids, as predicted by both PredAlgo and TargetP (Bagnato et al., 2017b). Contiguous to this fragment, the sequence features a region of about 223 amino acids with no apparent structure. In this disordered portion, we previously predicted at least two putative acyltransferase motifs that include the conserved histidine that most frequently acts as catalytic residue (Bagnato et al., 2017b). To the carboxy end, DGAT3 has a conserved 2Fe-2S cluster-binding domain, which we previously modeled on the thioredoxin-like ferredoxin of *Aquifex aeolicus* (Yeh et al., 2000), followed by a carboxy end with no significant similarity to any characterized motif.

**Figure 1.**
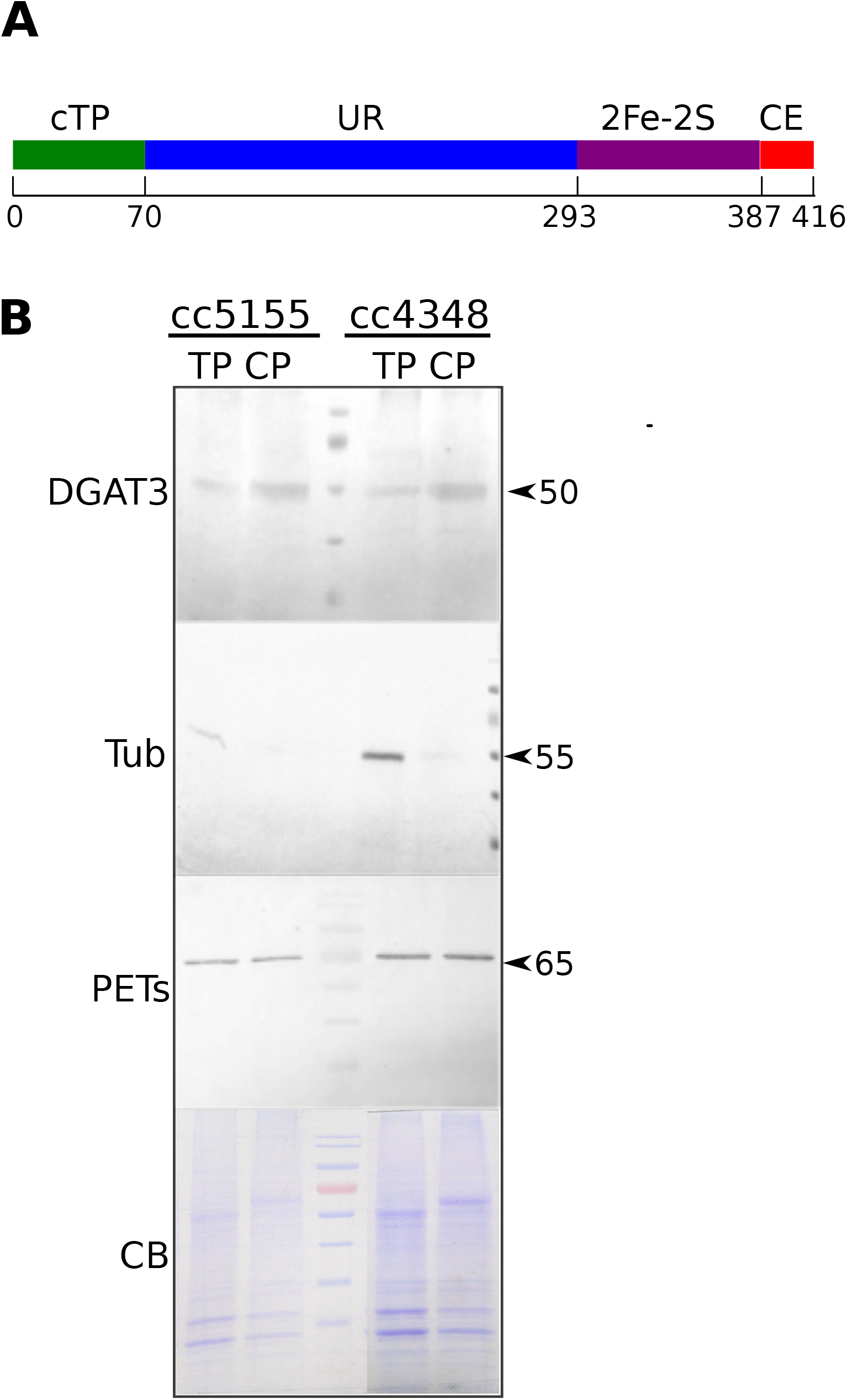
*Chlamydomonas reinhardtii* DGAT3 localizes to the chloroplast. **A)** Schematic representation of DGAT3 protein showing the predicted chloroplast transit peptide (cTP), the amino terminal unstructured region (UR), the 2Fe-2S cluster-binding domain (2Fe-2S) and the non-homologous carboxy end (CE). The numbers below are the predicted amino acid positions of each domain junction. **B)** Western blot analysis of total protein (TP) and chloroplast protein (CP) fractions from *C. reinhardtii* strains cc-5155 and cc-4348. Five micrograms of proteins per lane were separated in 12% SDS-polyacrilamide gels and either stained with Coommassie Brilliant Blue (CB) or analyzed by western blot using anti-DGAT3, anti-tubulin (Tub) and anti-PETs antibodies. Arrowheads indicate approximate molecular weights, estimated by comparison with molecular size standards.

In order to confirm the chloroplast targeting prediction, we isolated *C. reinhardtii* chloroplasts from two cell wall-less strains, cc-5155 and cc-4348 (*sta6*) and analyzed the presence of DGAT3 by western blot analysis. Figure 1B shows that DGAT3 protein was present in both total protein and chloroplast protein samples from both strains. As expected, the chloroplast polyprotein of elongation factor Ts (PETs) (Beligni et al., 2004) was also detected in both fractions, whereas only traces of beta tubulin, a cytosolic marker, were observed in chloroplasts. These results indicate that *C. reinhardtii* DGAT3 is indeed a chloroplast protein.

All proteins containing 2Fe-2S cluster-binding domains characterized so far act as low potential electron carriers (Fukuyama, 2004; Zu et al., 2002). This prompted us to hypothesize that DGAT3 might accomplish the synthesis of TAGs in conjunction with the photosynthetic machinery and, hence, respond to light. TAGs were previously shown to increase in *C. reinhardtii* cc-124 cells after transferring them from low light to saturating light (Goold et al., 2016), but the enzymes responsible for this accumulation remain unknown. With the purpose of analyzing a potential connection between DGAT3 and light, we evaluated the effect of shifting cells from dark to light in classical light-activation experiments (Beligni et al., 2004; Coragliotti et al., 2011). Cells were cultured in Tris-Acetate-Phosphate (TAP) media under continuous light of 4,500 lux until they reached a cell density of 3 × 10^6^ cells/ml, transferred to the dark for 16 h and then shifted back to the initial light condition (4,500 lux). This experiment was done with both wild type cc-125 and cc-4348 cells. The cc-4348 strain is cell wall-less and has a deletion in the *STA6* gene (encoding a subunit of ADP-glucose pyrophosphorylase), which renders this strain deficient in starch synthesis (Zabawinski et al., 2001). Expression was determined by RT-qPCR. Two genes were evaluated as potential endogenous controls, *actin* and *gblp* (Table 1). According to the analysis of RT-qPCR data, *gblp* levels resulted stable in the experiments tested (Supplementary Data, Tables S1 and S2), whereas *actin* was only relatively stable in cc-4348 experiments and fluctuated considerably upon transferring cc-125 cells from dark to light (data not shown). By virtue of these results, the expression levels of all target genes were subsequently normalized to *gblp* levels.

**Table 1.**
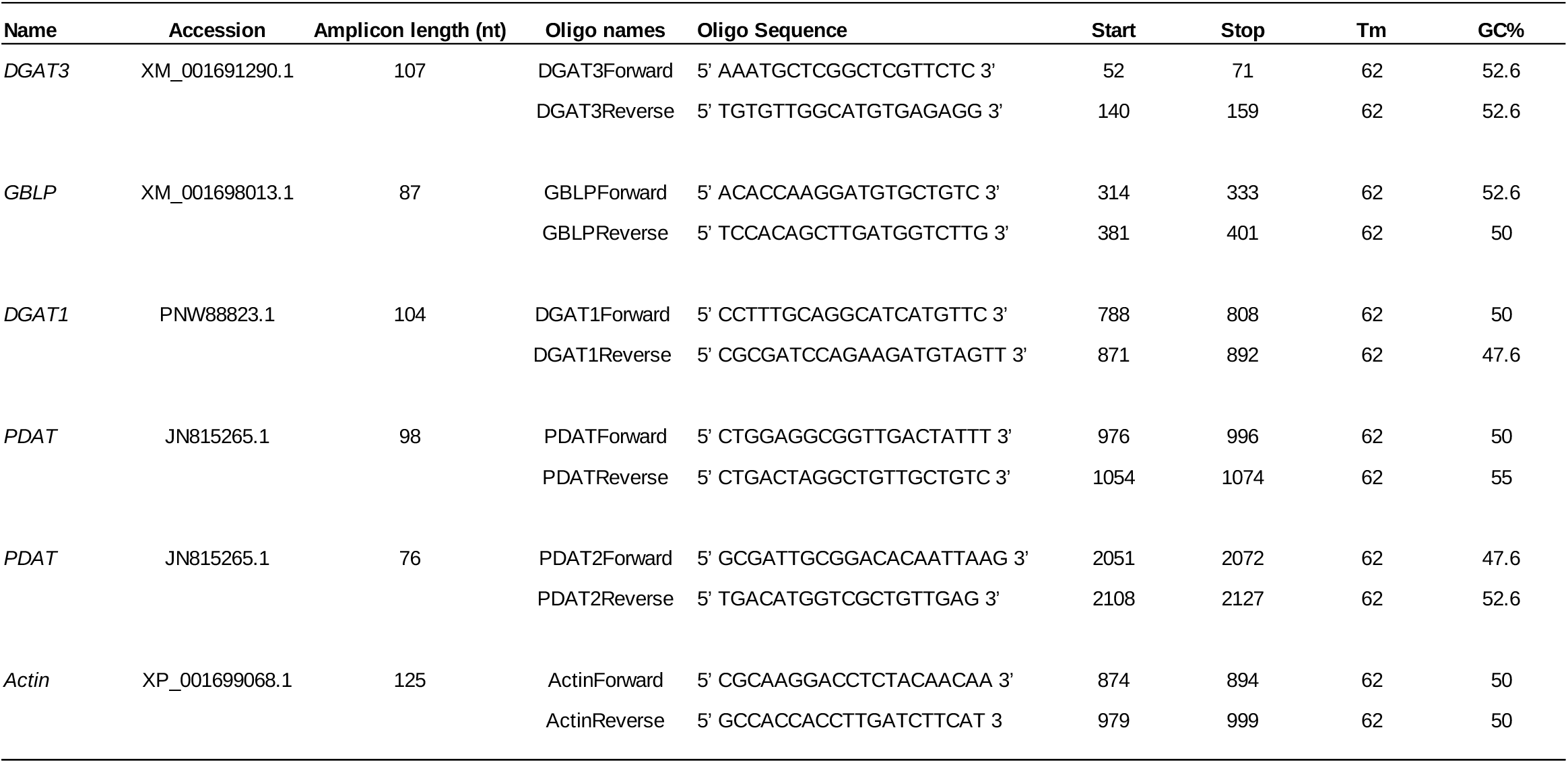
Summary of oligonucleotides used for RT-qPCR and amplicon position and length for each of the genes analyzed.

Figure 2A shows that, in the cc-4348 strain, *dgat3* mRNAs increased 2-fold as early as 1 h into the light period, peaked at about 12 h (an average value of about 4-fold) and remained elevated until the end of the experiment (3-fold). In cc-125 cells, no significant variations in *dgat3* mRNAs were observed between the end of the dark and the light period (Figure 2A, Table S3). In the same assays, TAGs accumulated in cc-4348 upon transferring the cells to light, reaching a maximum intensity at 6 h into the light period and decreasing considerably at 24 h (Figure 2B). In contrast, in cc-125 cells, no remarkable variations in TAG content were observed during the first 12 h of light compared to the end of the dark, and only a decline at 24 h could be observed (Figure 2B, Figure S1). These results show that DGAT3 expression and TAG content behave similarly upon shifting TAP-cultured cells from dark to moderate light conditions. While both *dgat3* mRNA levels and TAGs increased within the first 12 h into the light period in the cc-4348 strain, they remained fairly stable in the wild type strain.

**Figure 2.**
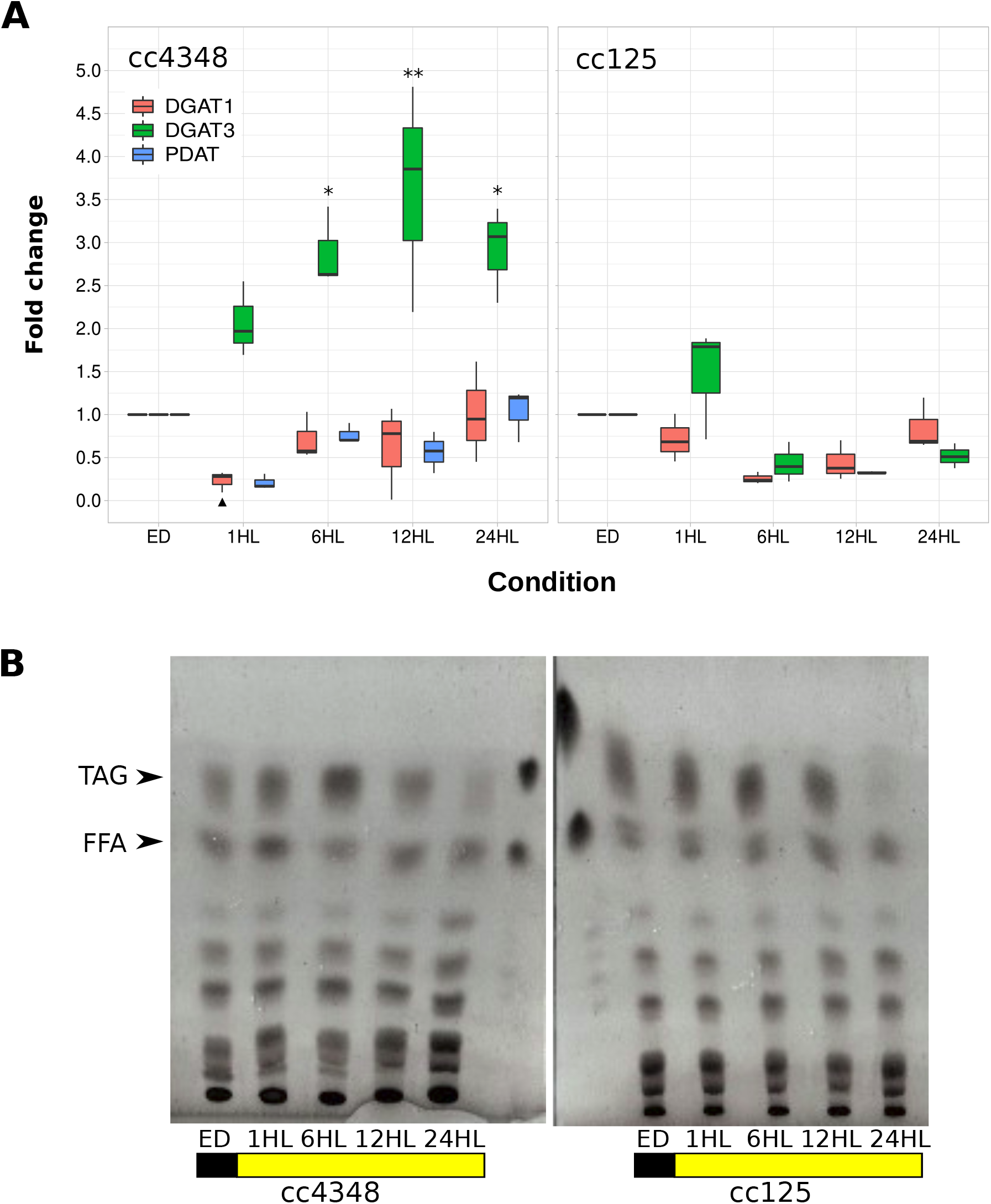
Exposure to moderate light triggers accumulation of DGAT3 mRNAs and TAGs in *sta6* cells in the presence of acetate. *C. reinhardtii* cells from wild type (cc-125) and the *sta6* mutant (cc-4348) were grown in TAP media under continuous light (4,500 lux) to approximately 3 × 10^6^ cells/ml, transferred to dark conditions for 16 h and re-transferred to 4,500 lux of light. Samples were harvested at the end of the dark period (ED) and 1 h, 6 h, 12 h and 24 h after shifting the cells to light (1HL, 6HL, 12HL and 24HL, respectively). **A)** Box plots showing the expression of *dgat3, dgat1* and *pdat* mRNAs analyzed by RT-qPCR. Results are shown as the fold change of each gene normalized to the endogenous control (*gblp*) in the same condition and to the ED sample. Values for independent biological triplicates are shown, vertical bars indicate minimum and maximum values, horizontal black strips indicate median values. Statistical analyses were done using dN0 values by means of one-way ANOVA and Dunnett’s multiple comparisons test; significant differences of samples against the ED sample are shown, ** P < 0.01, * P < 0.05, ▲ P < 0.1. **B)** Thin layer chromatography (TLC) of lipids extracted using the Bligh and Dyer method, separated in silicagel G-60 plates using hexane:sulphuric ether:formic acid (80:20:2, v/v) as solvent system and developed by charring. Sample volumes equivalent to 12 million cells were loaded on each lane. Arrowheads show the spots of triacylglycerol (TAG) and free fatty acid (FFA) standards. The black box represents the dark period, the yellow box represents moderate light. A representative image of three independent biological replicates is shown.

To determine the response to light of other enzymes that perform TAG synthesis, we analyzed the mRNA levels of DGAT1 and phospholipid diacylglycerol acyltransferase (PDAT). We selected those two enzymes since both have the potential to carry out TAG synthesis in the chloroplast, according to previous evidence (Bagnato et al., 2017b; Yoon et al., 2012). Quality control of RT-qPCR data indicated that the results obtained for *dgat1* mRNAs were sound, whereas *pdat* RT-qPCRs were only reliable for cc-4348. For this last strain, although the high counts (Cts) obtained suggested that the levels of *pdat* mRNAs were considerably low, the melting curves and agarose gels obtained showed that a single, pure amplicon was being amplified (Tables S1 and S2). For cc-125 *pdat* mRNAs, Cts were even higher and the melting curves contained multiple peaks, turning the analysis unreliable (data not shown).

Figure 2A shows that both *dgat1* and *pdat* mRNAs decreased moderately in TAP-grown cc-4348 cells 1 h after shifting the cells from dark to light and returned to the initial levels between 6 and 24 h into the light period. In cc-125 cells, no significant variations in *dgat1* mRNAs were observed between the end of the dark and the light period (Figure 2A, Table S3). In general, these results suggest that TAG accumulation induced by moderate light in cc-4348 cells cannot be explained by the pattern of *dgat1* and *pdat* mRNAs in our experimental conditions.

TAG accumulation in the chloroplast has been reported to occur only in the presence of acetate (Fan et al., 2011; Goodenough et al., 2014; Goodson et al., 2011). With the aim of analyzing if this pattern also held true for DGAT3 induction in cc-4348 cells, we analyzed *dgat3* mRNAs and TAG accumulation in this strain cultured in minimal media (without acetate) and transferred from dark to light as in the previous experiment. Figure 3A shows that *dgat3* mRNAs were reduced significantly upon transferring the cells from dark to light in minimal media. In the same conditions, TAG content remained relatively stable for the initial hours after shifting the cells to light and decreased mildly at 24 h (Figure 3B, Figure S1). Hence, we concluded that both light-activated DGAT3 expression and TAG accumulation depend upon the presence of acetate in the media in cc-4348 cells, in our experimental conditions.

**Figure 3.**
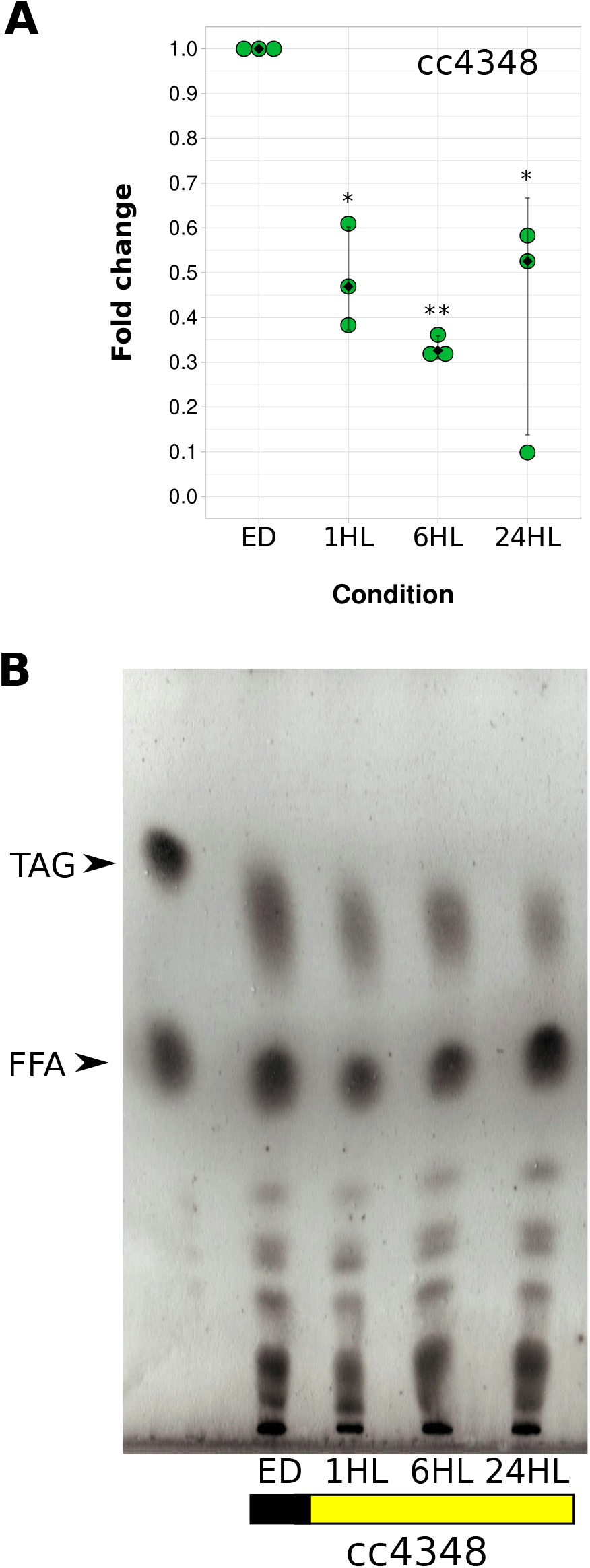
Response of DGAT3 mRNAs and TAG levels in *sta6* cells grown in minimal media. Cells from the *sta6* mutant (cc-4348) were grown in Tris-minimal media under continuous light (4,500 lux) to approximately 3 × 10^6^ cells/ml, transferred to dark conditions for 16 h and re-transferred to 4,500 lux of light. Samples were harvested at the end of the dark period (ED) and 1 h, 6 h and 24 h after shifting cells to light (1HL, 6HL and 24HL, respectively). **A)** Dot plot showing the expression of *dgat3* mRNAs analyzed by RT-qPCR. Results are expressed as the fold change of *dgat3* normalized to the endogenous control (*gblp*) in the same condition and to the ED sample. Green circular dots show the values of independent biological replicates. Vertical bars indicate standard deviations, black diamond dots show median values. Statistical analyses were done using dN0 values by means of one-way ANOVA and Dunnett’s multiple comparisons test; significant differences against the ED sample are shown, ** P < 0.01, * P < 0.05. **B)** Thin layer chromatography (TLC) of lipids extracted using the Bligh and Dyer method, separated in silicagel G-60 plates using hexane:sulphuric ether:formic acid (80:20:2, v/v) as solvent system and developed by charring. Sample volumes equivalent to 30 million cells were loaded on each lane. Arrowheads show the spots of triacylglycerol (TAG) and free fatty acid (FFA) standards. The black box represents the dark period, the yellow box represents moderate light. A representative image of three independent biological replicates is shown.

The differences in the accumulation of *dgat3* mRNAs and TAGs between cc-125 and cc-4348 cells shown in Figure 2 are most likely due to cc-4348 deficiency in starch accumulation. Light provides the means to generate reducing power that needs to be harnessed properly in order to protect the photosystems. The relatively high concentration of acetate in TAP media imposes the necessity to store the excess of carbon. In cc-4348, TAGs are the main storage molecules for both carbon and reducing power and, hence, it is not surprising that their levels increase upon transferring the cells from dark to moderate light (4,500 lux). We hypothesized that we could achieve a similar effect in cc-125 cells by increasing the light intensity, in order to subject the cells to a condition in which the excess of reducing power could not be effectively stored solely into starch. To this end, we measured *dgat3* mRNA levels and TAG accumulation in cc-125 cells cultured in TAP media at 4,500 lux, transferred to the dark for 16 h and then shifted to high light (15,000 lux). Figure 4A shows that, unexpectedly, *dgat3* mRNAs declined upon transferring cc-125 cells from dark to high light. In the same conditions, TAGs remained relatively stable for the first 12 h, and declined at 24 h into the light (Figure 4B), similarly to the experiment in Figure 2B. When analyzing each sample into detail, we realized that the intensity of the TAG signals in the ED samples relative to the samples harvested in the light, at both moderate and high light, was substantially higher for cc-125 (Figures 2B and 4B) than for the cc-4348 strain (Figure 2B). Since the levels of TAGs after a prolonged period at 4,500 lux were relatively low in the cc-125 strain (Figures 2B and 4B, 24HL samples), we assumed that the high levels of TAGs at the ED were originated from dark accumulation, probably due to a reduction of O_2_ in non-aerated, batch cultures, as reported previously (Hemschemeier et al., 2013). In agreement with this assumption, we observed that TAG levels increased during a 48 h-dark period and decreased slowly during the subsequent light period in cc-125 cells (Figure S1).

**Figure 4.**
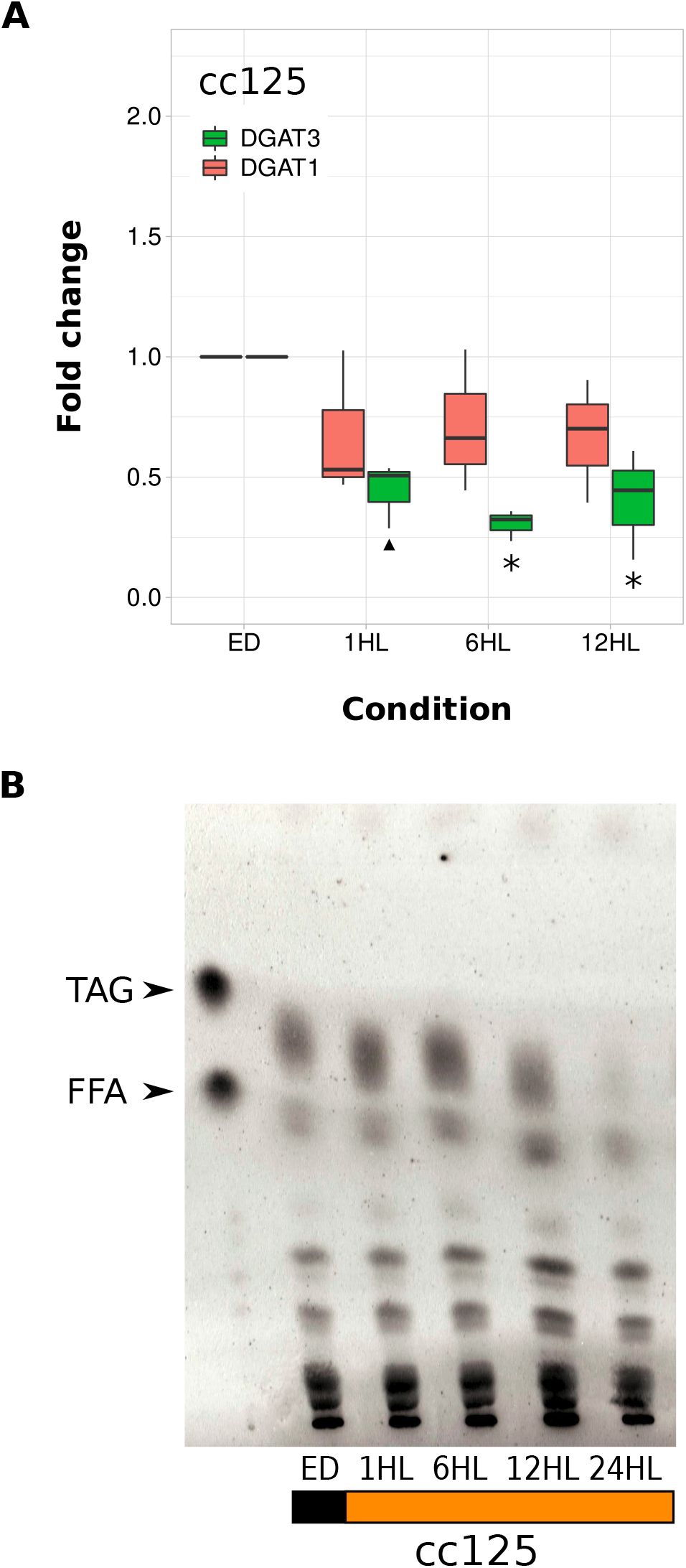
DGAT3 mRNAs and TAGs decrease in the wild type strain after shifting cells from darkness to high light. Cells from cc-125 were grown in TAP media under continuous light (4,500 lux) to approximately 3 × 10^6^ cells/ml, transferred to dark conditions for 16 h and then shifted to 15,000 lux of light. Samples were harvested at the end of the dark period (ED) and 1 h, 6 h, 12 h and 24 h after shifting cells to light (1HL, 6HL, 12HL and 24HL, respectively). **A)** Box plots showing the expression of *dgat3* and *dgat1* mRNAs analyzed by RT-qPCR. Results are expressed as the fold change of each gene normalized to the endogenous control (*gblp*) in the same condition and to the ED sample. Values for independent biological triplicates are shown, vertical bars indicate minimum and maximum values, horizontal black strips indicate median values. Statistical analyses were done using dN0 values by means of one-way ANOVA and Dunnett’s multiple comparisons test; significant differences against the ED sample are shown, * P < 0.05, ▲P < 0.1. **B)** Thin layer chromatography (TLC) of lipids extracted using the Bligh and Dyer method, separated in silicagel G-60 plates using hexane:sulphuric ether:formic acid (80:20:2, v/v) as solvent system and developed by charring. Sample volumes equivalent to 10 million cells were loaded on each lane. Arrowheads show the spots of triacylglycerol (TAG) and free fatty acid (FFA) standards. The black box represents the dark period, the orange box represents high light. A representative image of three independent biological replicates is shown.

In order to evaluate the response to light without subjecting cc-125 cells to a dark period, we evaluated DGAT3 and TAG content by replacing the dark interval with a low light (1,500 lux) period of the same length, after which cells were transferred to high light. Figure 5A shows that *dgat3* mRNAs increased 1 h after transferring the cells to high light, peaked at 6 h into the high light period (ca. 5-fold) and declined at longer times, remaining at about 2.5-fold until the end of the experiment. In this case, *dgat1* mRNAs also showed a moderate but steady increase of approximately 2-fold between 1 and 12 h into the high light period, decreasing again to the initial levels at the end of the experiment. Figure 5B shows that, at the end of the low-light period (ELL), TAG levels were relatively low and increased 1 h into the high light period, returning to basal levels afterwards. Altogether, these results show that induction of DGAT3 expression and TAGs by light also occurs in the cc-125 strain.

**Figure 5.**
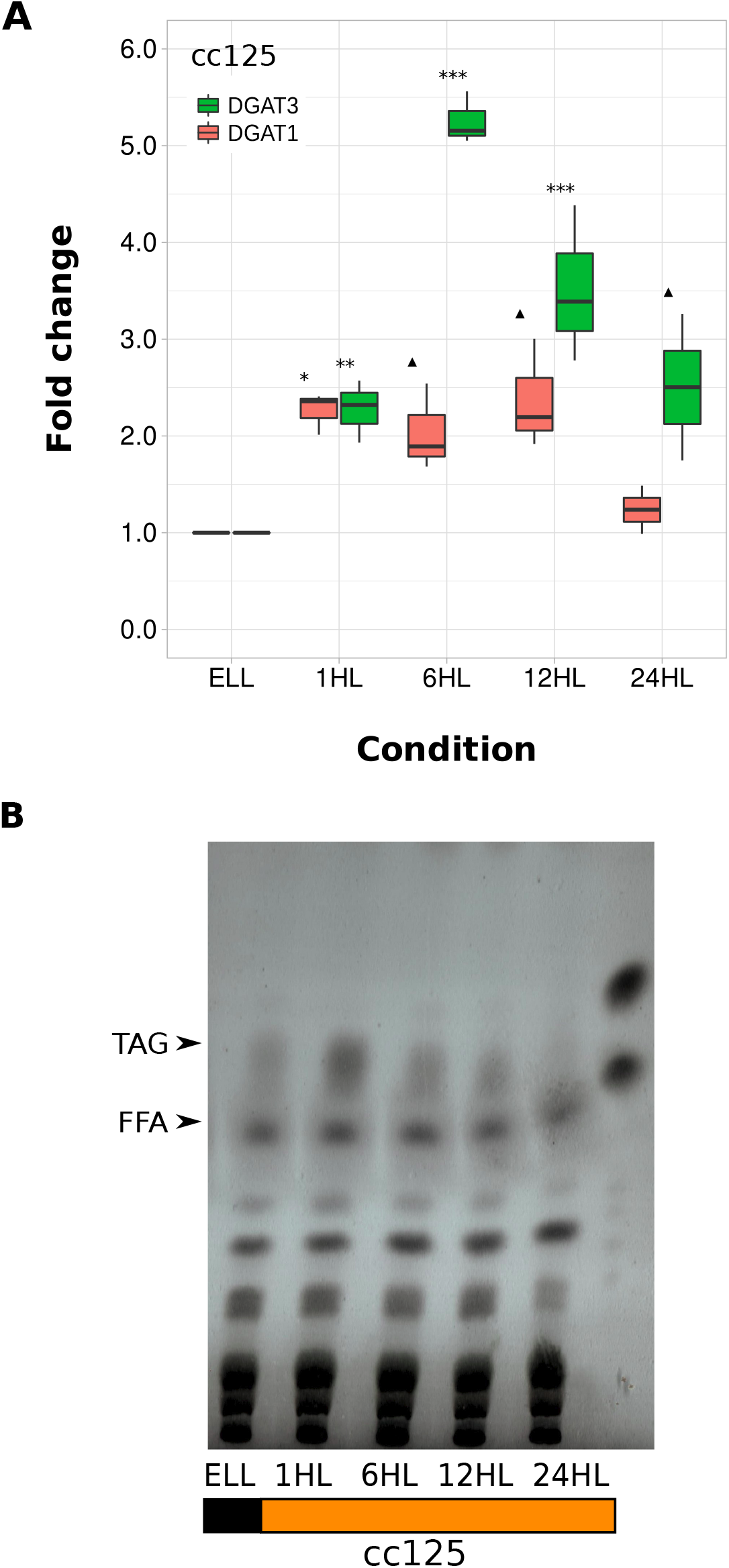
DGAT3 mRNAs and TAGs accumulate in the wild type strain after shifting cells from low light to high light. Cells from the wild type cc-125 strain were grown in TAP media under continuous light (4,500 lux) to approximately 3 × 10^6^ cells/ml, transferred to 1,500 lux for 16 h and then switched to 15,000 lux of light for 24 h. Samples were harvested at the end of the low light period (ELL) and 1 h, 6 h, 12 h and 24 h after shifting cells to high light (1HL, 6HL, 12HL and 24HL, respectively). **A)** Box plots showing the expression of *dgat3* and *dgat1* mRNAs analyzed by RT-qPCR. Results are expressed as the fold change of each gene in each condition compared to the endogenous control (gblp) in the same condition. Vertical bars indicate minimum and maximum values, horizontal black strips indicate median values. Statistical analyses were done using dN0 values by means of one-way ANOVA and Dunnett’s multiple comparisons test; significant differences against the ED sample are shown, *** P < 0.001, ** P < 0.01, * P < 0.05, ▲ P < 0.1. **B)** Thin layer chromatography (TLC) of lipids extracted using the Bligh and Dyer method, separated in silicagel G-60 plates using hexane:sulphuric eter:formic acid (80:20:2, v/v) as solvent system and developed by charring. Sample volumes equivalent to 10 million cells were loaded on each lane. Arrowheads show the spots of triacylglycerol (TAG) and free fatty acid (FFA) standards. The black box represents the dark period, the orange box represents high light. A representative image of three independent biological replicates is shown.

## Discussion

In this work, we report that light-induced TAG accumulation in *C. reinhardtii* occurs in agreement with the expression of DGAT3, a soluble, chloroplast-localized DGAT. In the conditions investigated, TAG production could not be fully explained by changes in the expression of DGAT1 or PDAT, two enzymes that could potentially synthesize TAGs on the chloroplast envelope. These results point to DGAT3 as a feasible and exciting candidate for light-activated TAG synthesis in the chloroplast, as shown in the model of Figure 6.

**Figure 6.**
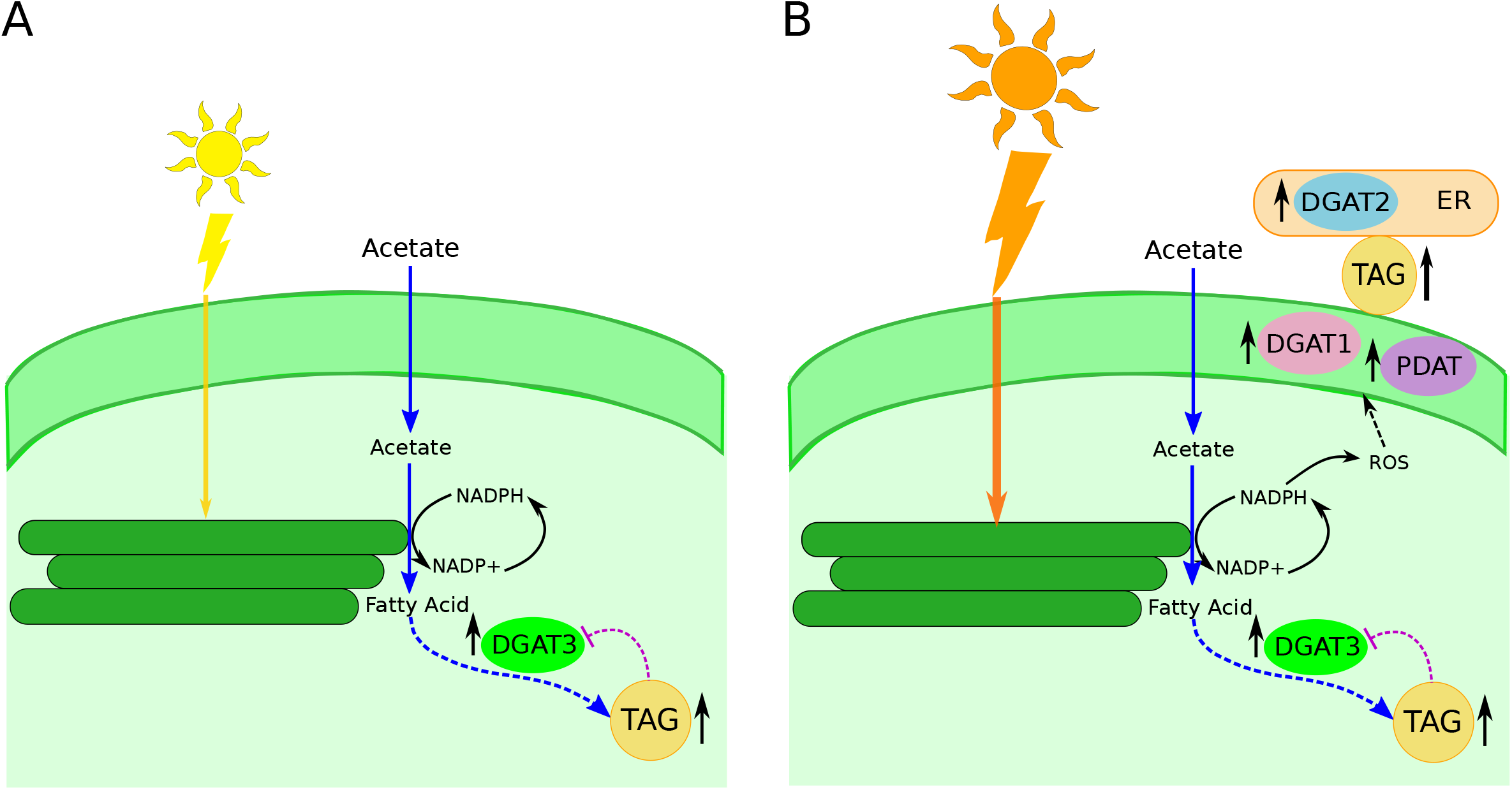
Schematic model of light-induced TAG accumulation in *C. reinhardtii.* **A)** When cells are transferred from dark to limiting light conditions in the presence of acetate, a pulse increase in DGAT3 expression occurs (upward black arrow by DGAT3). We hypothesize that reducing power provided by light and organic carbon supplied in the form of acetate are directed, as part of the regulation of this enzyme, towards the synthesis of TAGs in the chloroplast, which also increase with moderate light (upward black arrow by TAG). When the levels of TAGs reach a certain level, we propose that a regulatory loop inhibits further DGAT3 expression and TAG synthesis (purple dashed line). This mechanism could prevent light activation in cc-125 cells that come from a hypoxic dark period and, thus, have elevated pre-existing TAG levels. **B)** When cells are transferred to high light, we hypothesize that DGAT3-mediated TAG accumulation is not enough to harness the electron excess furnished by light. Hence, the expression of stress/ROS-related enzymes, such as DGAT1 (this work), PDAT and DGAT2 (Chouhan et al., 2020) is activated. We cannot conclude from our results what the relative contribution of each enzyme under high light is or whether DGAT3 expression is activated at irradiances other than those evaluated in this work, but we propose that the responses shown in B) are activated gradually as more photons that are absorbed cannot be stored by the mechanisms in A). In addition, while light response to limiting light might only involve the chloroplast, the response to high light might occur in both the plastid and the ER/cytosol, as proposed previously (Chouhan et al., 2020; Goold et al., 2016). Dashed lines correspond to processes yet to be fully demonstrated. Only the synthesis of TAGs is shown, without addressing their final storage.

In algae, TAG accumulation is mostly associated with the canonical ER/cytosol pathway (Goold et al., 2016; Tsai et al., 2015; Weng et al., 2014). However, under certain conditions, TAGs have also been proposed to accumulate in association with the chloroplast (Fan et al., 2011; Goodson et al., 2011). Goold et al (2016) reported that saturating light triggers TAG synthesis in *C. reinhardtii,* both inside and outside the plastid, mostly associated with the chloroplast envelop. In this work, we report that DGAT3 expression and TAG accumulation do not only respond to high light, but undergo light-dependent activation in a broad sense, since their levels also increase at moderate light intensities, as shown in Figure 6A. True light activation is expected to operate to some extent at all light intensities, including limiting light conditions, in which the rate of light absorption is well matched with the rate of photosynthesis (Björkman and Demmig, 1987). In our case, an increment in the content of TAGs after the transition from dark to limiting light (4,500 lux) occurred in the *sta6* mutant and matched DGAT3 induction. Neither of the other two enzymes evaluated (DGAT1 and PDAT) could explain the boost of TAGs in limiting light in the *sta6* mutant. The evaluation of those two enzymes was not circumstantial. PDAT is targeted to the chloroplast in *C. reinhardtii* and can synthesize TAGs from several substrates (Yoon et al., 2012). DGAT1, though frequently assumed to be localized to the ER, has a predicted chloroplast transit peptide and could be anchored to the chloroplast envelope (Bagnato et al., 2017b). Hence, although we cannot rule out that either post-translational activation or the expression other enzymes (e.g. DGAT2, phytyl ester synthases) participate in moderate light-activated TAG production, the strong agreement between TAG levels and DGAT3 expression provides the appealing possibility that the accumulation of TAGs in response to limiting light is catalyzed by DGAT3. Unfortunately, there are no DGAT3 mutants available in the Chlamydomonas Library Project (CLiP), but future work will be aimed at following a reverse genetics approach to interrogate the function of the *dgat3* gene.

The effect of high light, although related, has significant differences with light activation. In high light, it has been demonstrated that the rate of photosynthesis becomes saturated, but light in excess is still absorbed and can damage the photosystems and other macromolecules within the cell (Niyogi, 1999). Hence, the response to high light is probably a combination of light activation and stimulation of photoprotective mechanisms. In our case, TAG accumulation in wild type cc-125 cells at high light (15,000 lux) was accompanied by an up-regulation of both *dgat3* and *dgat1* mRNAs. Even though not all enzymes that can synthesize TAGs have been evaluated in our high light experiments, one could hypothesize that a more general response to stress is activated in these conditions compared to moderate light. Recently, it was demonstrated that high light induces ROS, which leads to TAG accumulation, in agreement with an increase in DGAT2A and PDAT (Chouhan et al., 2020). All these results suggest that light activation and light stress operate via related, but not necessarily identical, pathways. We propose that, while limiting light might only activate DGAT3 expression and TAG accumulation in the chloroplast (Figure 6A), high light might also activate the expression of other enzymes that participate either in the chloroplast or the ER, as shown in Figure 6B and as also proposed previously (Chouhan et al., 2020; Goold et al., 2016). In addition, while DGAT3 expression might be a target of the light signal transduction cascade, other enzymes involved in TAG synthesis might be targets of stress/ROS signaling cascades (Figures 6A and 6B). It is noteworthy that, in both limiting light and high light, the increase in *dgat3* mRNAs and TAG levels occurred as a pulse shortly after transferring the cells to light, consistent with light activation, as has been shown for other targets (Beligni et al., 2004; Gray et al., 2003; Idoine et al., 2014; Salvador et al., 1993). DGAT3 mRNAs were previously shown to also respond to nitrogen deprivation, but this occurred only at long periods after imposing this stress (Boyle et al., 2012; Goodenough et al., 2014; Liu et al., 2016). In the same N deplete conditions, DGAT1, PDAT and a few of the DGAT2 isoforms had a more rapid response. Altogether, these results suggest that DGAT1, DGAT2 and PDAT are mostly related to stress as already proposed, whereas DGAT3 might be mainly regulated by light and, perhaps indirectly or circumstantially, moderately responsive to sources of stress such as nitrogen deprivation.

The idea that algae accumulate TAGs as a sink for excess photosynthetic energy and reductant in order to prevent photochemical damage was proposed long ago (Roessler, 1990). In this widely accepted hypothesis (Du and Benning, 2016; Hu et al., 2008; Klok et al., 2014; Li et al., 2012; Solovchenko, 2012), photosynthetic carbon and energy assimilation that can no longer be directed to growth result in the storage of excess reduced carbon (overflow products), both in the form of starch and TAGs. In this scenario, the differences in the response of DGAT3 and TAGs to light between the wild type and *sta6* strains could be explained by the fact that *sta6* cells cannot synthesize starch and have, hence, an increased pressure for the storage of reduced carbon in the form of TAGs. In this work, we provide evidence that advances further into the differences between the two types of metabolism.

Our TLCs showed that, in wild type cells, TAGs increased in the dark in batch cultures and remained high until the end of this period, whereas *sta6* cells had low levels of TAGs at the end of the dark. As Hemschemeier et al (2013) previously reported, TAGs accumulate in *Chlamydomonas* during extended darkness (24 h) in TAP media, due to the reduction of oxygen concentration in batch cultures. In our experiments, wild type cells incubated in the dark most likely generated energy and reducing power from acetate consumption and starch degradation, with TAG levels rising due to hypoxia. In contrast, in *sta6* cells, the effect of oxygen reduction on TAG accumulation was evidently counteracted by the need to catabolize TAGs for energy and reducing power in the dark. We propose that the differences in light activation between wild type and *sta6* cells are related to this. After a 16-h period of darkness, neither DGAT3 expression nor TAGs increased upon transferring wild type cells to light, and we hypothesize that this is related to the high levels of TAGs at the end of the dark, since the situation changed when darkness was replaced by low light. At the end of a 16 h low-light period, the levels of TAGs were low, more similar to the levels in the *sta6* strain at the end of the dark. In this condition, a pulse increase in DGAT3 and TAGs could be observed after shifting the cells to high light, evidencing that DGAT3 and TAGs do respond to light in the wild type. More broadly, our results suggest that light-activated DGAT3 expression and, hence, TAG accumulation are tied to the preexisting levels of TAGs. In other words, light-activated DGAT3 expression might be subjected to negative feedback inhibition by the end product TAG, as shown in the model of Figure 6. The existence of regulatory loops for TAG biosynthesis in *C. reinhardtii* has already been proposed (Liu et al., 2016). Since the over-accumulation of TAGs could lead to cell obesity, growth arrest and cell death, at least during nitrogen deprivation (Goodson et al., 2011), maintaining TAG levels at bay could ensure that the cell can properly store overflow products in a sustainable manner during the growth phase.

It was previously reported that the production of TAGs that occurs under nitrogen deprivation was dependent on exogenous acetate for both *C. reinhardtii cw15* and *sta6* strains (Goodenough et al., 2014; Goodson et al., 2011). In this work, the increase in TAGs and DGAT3 levels also depended on the presence of acetate after transferring *sta6* cells from dark to light. In minimal media, *dgat3* mRNA levels markedly decreased in the light, whereas TAG levels remained relatively stable for the first 12 h into the light period and decreased moderately at 24 h. These results extend the observations presented above and suggest that DGAT3 regulation, and eventually TAG synthesis, are at a crossroads between the provision of reducing power by light and carbon supply. The sequence features of DGAT3 are ideal for this type of control. The presence of a 2Fe-2S cluster-binding domain indicates that this protein could accept electrons, directly or indirectly, from the photosynthetic machinery. We could hypothesize that, as part of DGAT3 regulation, these electrons are further funneled towards the synthesis of fatty acids and their acylation onto molecules of DAG (Bagnato et al., 2017b). In this scenario, it is not surprising that the expression of this enzyme is regulated by the two key components of this process: light and carbon.

Although we propose that DGAT3 can synthesize TAGs in the chloroplast, we cannot conclude from our results the final destination of those TAGs. Chloroplast TAGs have been connected to chloroplast lipid droplets (LDs) (Goodenough et al., 2014; Goodson et al., 2011; Goold et al., 2016), but the existence of that entity has recently been questioned (Moriyama et al., 2018). Beyond this controversy, several reports have proposed the appealing idea that a direct metabolic flow occurs between the chloroplast envelope and cytosolic LDs, without the essential intervention of the ER (Moriyama et al., 2018; Goodenough et al., 2014; Liu and Benning, 2013). In addition, chloroplast TAGs could be stored into plastoglobules, lipo-protein structures intimately related to the thylakoid membranes that contain TAGs, among other lipophilic compounds (Steinmüller and Tevini, 1985; Tevini and Steinmüller, 1985). If we consider all these possibilities, DGAT3-dependent TAG synthesis could result in the accumulation of TAGs stored into either plastoglobules or LDs, the latter either in the chloroplast or the cytosol. Future research will surely shed light into the participation of this unique enzyme in the complex metabolism of TAGs in green algae.

## Materials and Methods

### Chlamydomonas reinhardtii strains and culture conditions

The wild type *C. reinhardtii* strain cc-125 (*137c, mt+ nit1 nit2*), the cell wall-less strain cc-5155 (*mt+ cw15)* and the cell wall-less, starch deficient strain cc-4348 *sta6* (*mt+ sta6-1*), which contains a deletion of the gene encoding an ADP-glucose pyrophosphorylase subunit and, hence, is incapable of starch formation (Zabawinski et al., 2001) were obtained from the *Chlamydomonas* Resource Center, University of Minnesota, MN, USA. Cells were maintained on Tris-Acetate-Phosphate (TAP) (Gorman and Levine, 1965) agar plates. Prior to experiments, cells were inoculated at 10^4^ cells mL^−1^ in 100 ml Erlenmeyer flasks containing either liquid TAP or Tris-minimal media (Gorman and Levine, 1965) and grown on a rotary shaker at 23°C and continuous white light (6,400K) at 4,500 lux (approximately 60 μmol photons m^−2^ s^−1^). TAP and Tris-minimal media have identical recipes, with the exception that TAP contains 18.3 mM acetate, while Tris-minimal is acetate-free. All experiments were started when cell densities were approximately 3 × 10^6^ cells mL^−1^. Unless specified, in dark-light experiments cells were transferred to either dark (D) or low light (LL; 1,500 lux, approx. 20 μmol photons m^−2^ s^−1^) conditions for 16 h. Next, cultures were transferred to light of either 4,500 lux or 15,000 lux (high light; approx. 210 μmol photons m^−2^ s^−1^). Samples were taken at 1 h, 6 h, 12 h and 24 h into the light period. Since we wished to analyze the effect of light on mRNA and lipid accumulation, the cultures were grown in batch mode until the end of the dark period, but were switched to semi-continuous mode after transferring them to the light, in order to maintain cell densities constant and avoid an effect of differential cell shading between samples. Previously (Coragliotti et al., 2011), we had done this by centrifuging cells from all time points and resuspending them in fresh TAP to avoid a possible effect of nutrients coming from different volumes of fresh medium between samples. However, we modified this procedure since it was recently reported that centrifugation induces lipid production (Kato et al., 2019). The modified protocol consisted in adding, every hour, the volume of fresh medium necessary to revert the cell density to the value at the end of the dark, which we had previously determined that did not impact the accumulation of the selected target molecules at the times points analyzed (unpublished results). In addition to the maintenance of cell densities, each sample harvested accounted for 10% or less of the total culture volume, in order to avoid drastic changes in culture volume that could affect light penetration. At the end of the experiments, cells were harvested by centrifugation at 2,000 × g for 10 min, supernatants were discarded and pellets were flash-frozen in liquid N2 and stored at −80°C until further use. At least three independent, non simultaneous, experiments were done for each type of experimental design.

### Chloroplast isolation

Cells were harvested at 2 × 10^6^ cells mL^−1^, pelleted at 1,000 × g for 10 min and resuspended in chloroplast isolation buffer (0.3 M sorbitol, 5 mM MgCl_2_, 5 mM EGTA, 5 mM EDTA, 20 mM Hepes-KOH pH 8.0, 40 mM NaHCO_3_). Cells were re-pelleted and washed in chloroplast isolation buffer two more times. The final suspension was pipetted into an ice-cold N2 decompression chamber (Parr Model 4639, Parr Instrument Company, illinois, USA). The chamber was pressurized at 150 psi for 2 min and de-pressurized at about 3 mL min^−1^. The cells collected at the release tube were loaded onto a 20/45/65 % volumetric 3-step Percoll gradient. In such a gradient, intact cells will sediment at the bottom of the tube, whereas intact chloroplasts will sediment at the interphase between the 45 and 65 % layers and cell debris and cell contents will sediment in or on top of the 20 % layer. Percoll step solutions were prepared from a 95 % Percoll stock solution (95 % (v/v) Percoll, 3 % (w/v) PEG 6,000, 1 % (w/v) Ficoll and 1 % (w/v) BSA) diluted into a gradient mixture (25 mM Hepes-KOH pH 8.0, 10 mM EDTA, 5 % sorbitol). Gradients were centrifuged in a Sorvall HB-6 swinging bucket rotor at 6,000 × g for 20 min at 4 °C (with the brake off). Intact chloroplasts were carefully pipetted from the interphase, washed twice in chloroplast isolation buffer and resuspended in total protein lysis buffer for further analysis.

### RNA isolation and RT-qPCR

MIQE guidelines (Bustin et al., 2009) were taken into account for experimental design, sample manipulation and data analyses. RNAs were isolated as described previously (Coragliotti et al., 2011) with some modifications. The volumes equivalent to 10^7^ cells were harvested per sample and lysis was done in 1 mL final volume of 70 mM Tris-HCl pH 8, 200 mM NaCl, 20 mM EDTA, pH 8 and 2.7 % SDS at room temperature for 30 minutes. Homogenates were extracted twice with phenol: chloroform: isoamyl alcohol (25: 24: 1). The upper aqueous phases were precipitated with 1 volume of isopropyl alcohol overnight at −20°C. Pellets were washed with consecutive 70% and 100% ethanol washes and resuspended in diethyl pyrocarbonate (DEPC)-treated H_2_O once dry. RNAs were quantified in a Nanodrop (Thermo Scientific™ NanoDrop™ OneC Microvolume UV-Vis Spectrophotometer) by determining the Abs_260nm_ and purity was estimated with the Abs_260_/Abs_280_ and the Abs_260_/Abs_320_ ratios: only samples whose values were approximately 2.0 and 1.34, respectively, were used. In addition, RNA integrity was visualized in 0.9% agarose gels stained with Sybr™ Safe (Thermo Fisher Sci. Cat. #S33102). One microgram of total RNAs from each sample was treated with DNAse (Thermo Fisher Sci. Cat. #EN0521) and used for the synthesis of first strand cDNAs using MMLV (Thermo Fisher Sci. Cat. #28025013), both procedures according to the manufacturers’ instructions. Copy DNAs were stored at −20°C until further use. For RT-qPCR, *dgat3, dgat1, pdat, gblp* and *actin* mRNAs were analyzed. Details on GeneBank accession numbers, oligo sequences and length and position of the amplicons on the coding sequence of the mRNAs are shown in Table 1. The optimal temperature and dynamic range of each pair of oligos for each type of experiment were determined. We worked within template concentrations in which acceptable linearity (R2 ≥0.980) and efficiencies were observed. In addition, only comparisons between genes whose efficiencies differed by less than 10% were made (all between 100 and 109% efficiencies). Sybr-Green-based technology was used for RT-qPCR reactions; mixtures consisted in 2 μl of diluted template and 8 μl of FastStart Universal SYBR Green Master (Rox) (Cat. # 04913914001, Sigma-Aldrich Argentina). Reactions were set in technical duplicates or triplicates in MicroAmp® Fast Optical 48-well 0.1 mL reaction plates (Applied Biosystems, Cat. #4375816, through Invitrogen Argentina), covered with 48−well optical adhesive film (Applied Biosystems, Cat. #4375323, through Invitrogen Argentina) and run in an Applied Biosystems StepOne™ Real-Time PCR System with the following program: Holding stage: 95 °C for 10 min; cycling stage: 40 cycles of 95 °C for 15 sec (melting) and 60 °C for 1 min (annealing and extension), with an additional cycle for the determination of melting curves. Only conditions in which the reaction melting curve showed a single peak were analyzed, single amplicons were confirmed in agarose gels for selected samples. Data was analyzed using LinReg PCR (Ruijter et al., 2014) (Tables S1 and S2) to determine baselines and calculate individual PCR efficiencies for each sample. Starting concentrations for each sample, expressed in arbitrary fluorescence units (N0s) were used to calculate technical averages and to normalize target genes to reference genes and experimental conditions to control conditions.

### Protein extraction and western blot analyses

*C. reinhardtii* proteins were extracted in 50 mM Tris-HCl pH 8, 1 mM EDTA, 100 mM NaCl, 1 % SDS, 1 mM PMSF, 0.5 mM Leupeptin. Protein concentration in homogenates was determined with the bicinchoninic acid reagent using a bovine seroalbumin (BSA) standard curve. Samples were heated for 3 min at 95°C in 60 mM Tris-HCl pH 6.8, 12.5 mM EDTA, 2% SDS, 10% glycerol, 1% 2-mercaptoethanol and 0.02% bromophenol blue and separated in 12% SDS-polyacrilamide gels. PageRuler™ Plus prestained protein ladder (Thermo Fisher Sci. Cat. #26619, Invitrogen Argentina) was used as a size standard. Gels were either stained with 0.25% coomassie brilliant blue R-250 or transferred onto nitrocellulose for western blot analyses. Blots were blocked in 1x TSBT with 5% skimmed milk and incubated with primary antibodies overnight at 4°C on a rotary shaker. The primary antibodies used were: anti-*C. reinhardtii* DGAT3 (1:500 dilution of rabbit polyclonal antisera), anti-*C. reinhardtii* polyprotein of elongation factor Ts (PETs, 1:1000 dilution of rabbit polyclonal antisera) and mouse monoclonal anti-*Drosophila* beta-tubulin (E7 antibody, used at 0.5 μg/ml, obtained from the Developmental Studies Hybridoma Bank). E7 was deposited to the DSHB by Klymkowsky, M. (DSHB Hybridoma Product E7). Blots were washed in 1 × TBST and the corresponding secondary antibodies coupled to alkaline phosphatase were incubated at a 1:15,000 dilution for 1 h at room temperature. Blots were developed using nitroblue tetrazolium and 5-bromo-4-chloro-3-indolyl phosphate (Sigma Argentina).

### Lipid extraction and thin layer chromatography

Lipid extraction was done according to Bligh and Dyer (1959). Briefly, cells were resuspended in 3.8 ml of chloroform: methanol: H_2_O (1: 2: 0.8), vortexed for 20 sec and incubated at room temperature for 1 h, with occasional vortexing. Total lipid extraction was evidenced by the white color of the protein solid phase, which indicated that chlorophyll had completely partitioned to the chloroform phase. For phase partitioning, 1 mL chloroform and 1 mL H_2_O were added and tubes were centrifuged at 6,000 rpm for 2 min at 4 °C. The lower chloroform phase was transferred to another tube and extracts were dried for 1 h at 48 °C in a rotational vacuum concentrator (Martin Christ model RVC 2-18). Dried extracts were resuspended in a known volume of chloroform and spotted on 500 μm silica gel G-60 thin layer chromatography (TLC) glass plates. A mixture of hexane: sulphuric ether: formic acid (80: 20: 2, v/v) was used as running solvent system. Plates were run until the solvent front was 1 cm from the top of the plates (approx. 40 min) and dried for 1 h. Lipid spots were located after spraying the TLC plates with 3% CuSO4 (w/v), 8% H3PO 4 (v/v) and exposing them to 180°C for 10 min (Marsh and Weinstein, 1966). Lipid classes were identified by comparison to standards spotted along with the samples. Olive oil was used as a standard of TAGs. Oleic acid was used as free fatty acid standard and was kindly provided by Materia Hnos SACIF, Mar del Plata, Argentina (Product code MATSOL 101 OD). Biological replicates of TLCs shown in the main figures are shown in Figure S1.

### Data and statistical analyses

Graphs were done in RStudio, using the dplyr and ggplot2 packages (RStudio Team, 2020). For a better visualization of changes induced by light, RT-qPCR graphs were done using fold changes between the light conditions and the corresponding ED or ELL condition. For statistical analyses, dN0 values were used. These values are equivalent to dCt values, but corrected by LinRegPCR (Ruijter et al., 2014) for individual sample PCR efficiencies. Correlation between the data from each experiment was evaluated a using Pearson’s test. Residuals were calculated and used to test for normal distribution (https://home.ubalt.edu/ntsbarsh/Business-stat/otherapplets/Regression.htm). In the absence of evidence against normal distribution, one-way ANOVA was performed in RStudio. Significant differences between light conditions and the corresponding controls (ED or ELL) were evaluated using the post-hoc Dunnet’s test, using the RStudio DescTools package. A two-sided alternative hypothesis was used. Significant differences at p-values < 0.001, 0.01, 0.05 and 0.1 were reported. All the results of statistical analyses are shown in Table S3.

## Acknowledgements

We thank Materia Hnos SACIF, Mar del Plata, Argentina for their kind donation of oleic acid. This work was supported by funds to MVB from the Argentinean Consejo Nacional de Investigaciones Científicas y Tecnológicas (CONICET-PIP 11420110100090) and from Agencia Nacional de Promoción Científica y Tecnológica (ANPCyT, PICT-2013-2122). MVB, CB, GG and LM are CONICET Researchers, DS is a CONICET post-doctoral fellow.

