## Supplemental Figures 1 for "Expression of *Chlamydomonas reinhardtii* chloroplast diacylglycerol acyltransferase-3 is activated by light in concert with triacylglycerol accumulation"

**Supplementary Figure 1.** Additional biological replicates of the TLCs corresponding to the experiments shown in Figures 2-5 of the main manuscript, as well as one additional experiment. Lipids were extracted and TLCs were run as indicated in Materials and Methods. Experimental conditions and samples are indicated in each panel.

**A.** Strain cc-125, TAP media, dark to 4,500 lux of light. Sample volumes corresponding to 10 million cells per lane were loaded.

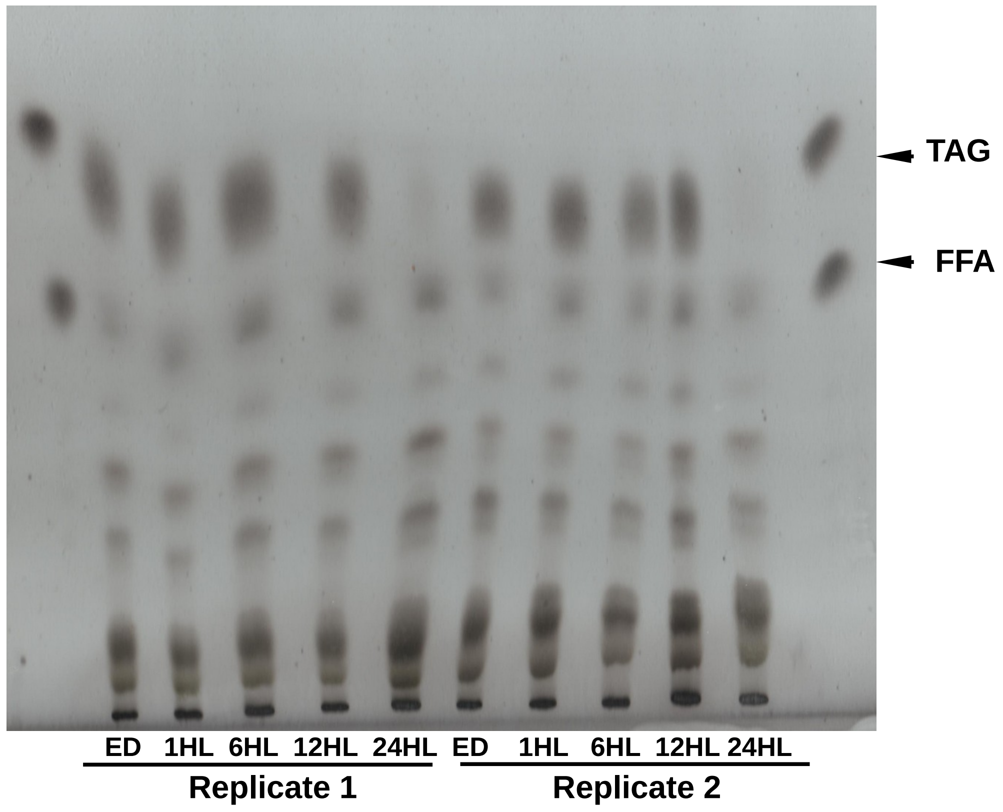


**B.** Strain cc-4348, TAP media, dark to 4,500 lux of light. Sample volumes corresponding to 5 (left panel and right panel) and 10 million cells (middle panel) per lane were loaded.

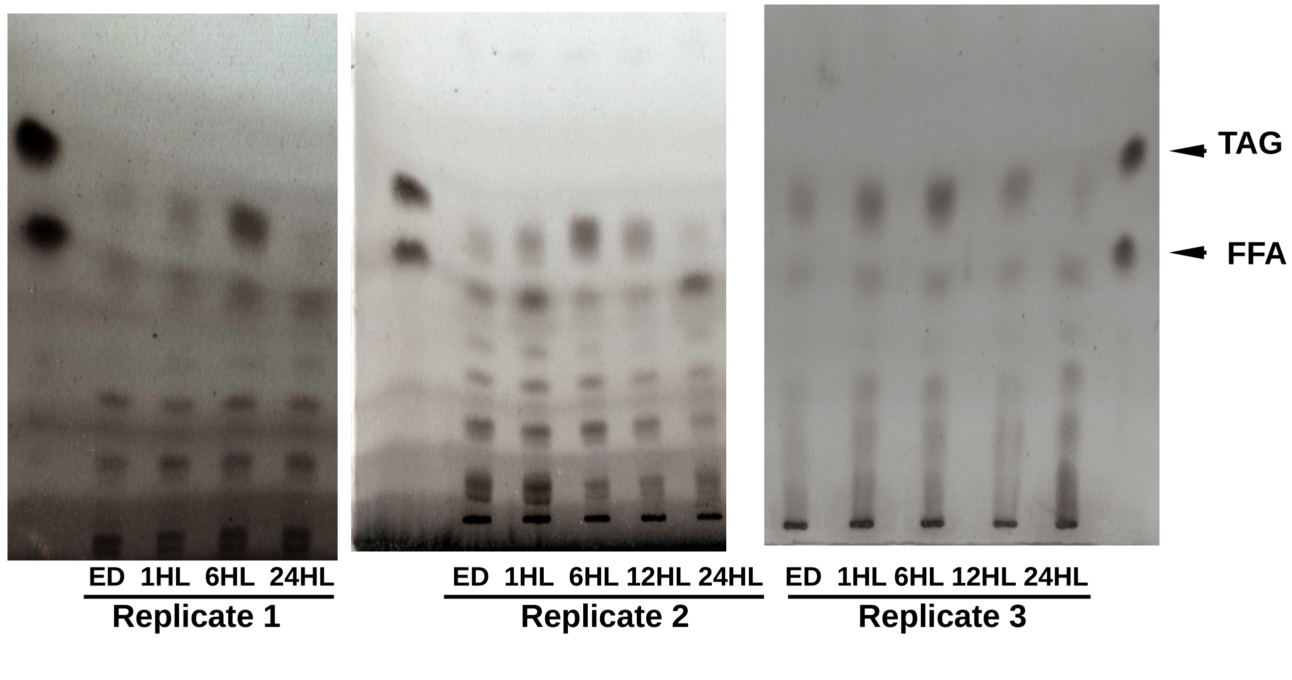

**C.** Strain cc-4348, Tris-minimal media, dark to 4,500 lux of light. Sample volumes corresponding to 25 million (Replicates 1 and 2) and 40 million cells per lane (Replicate 3) were loaded.

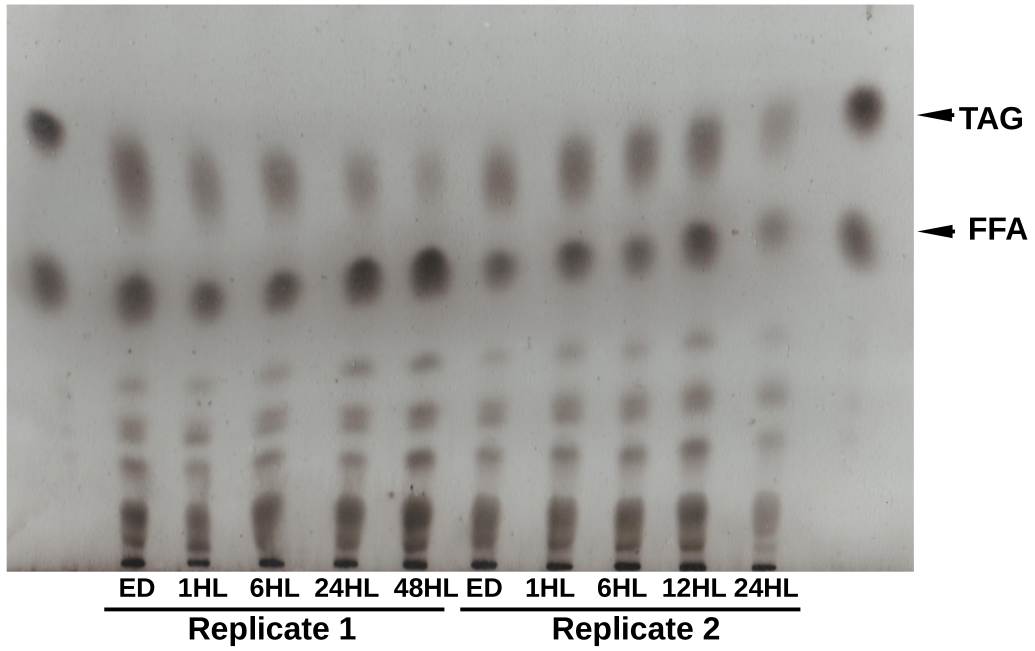


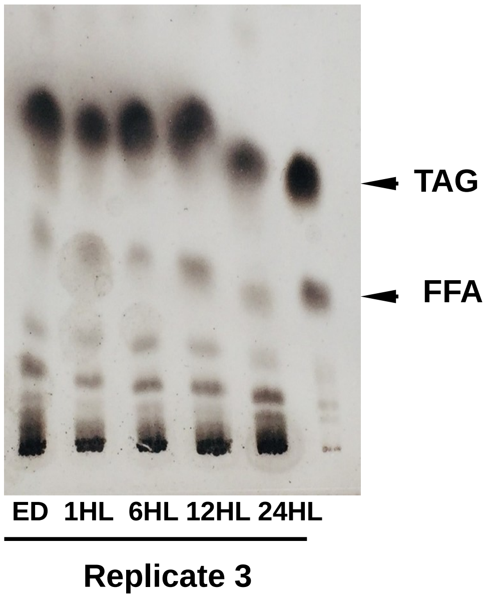


**D.** Strain cc-125, TAP media, dark to 15,000 lux of light. Sample volumes corresponding to 10 million cells per lane were loaded.

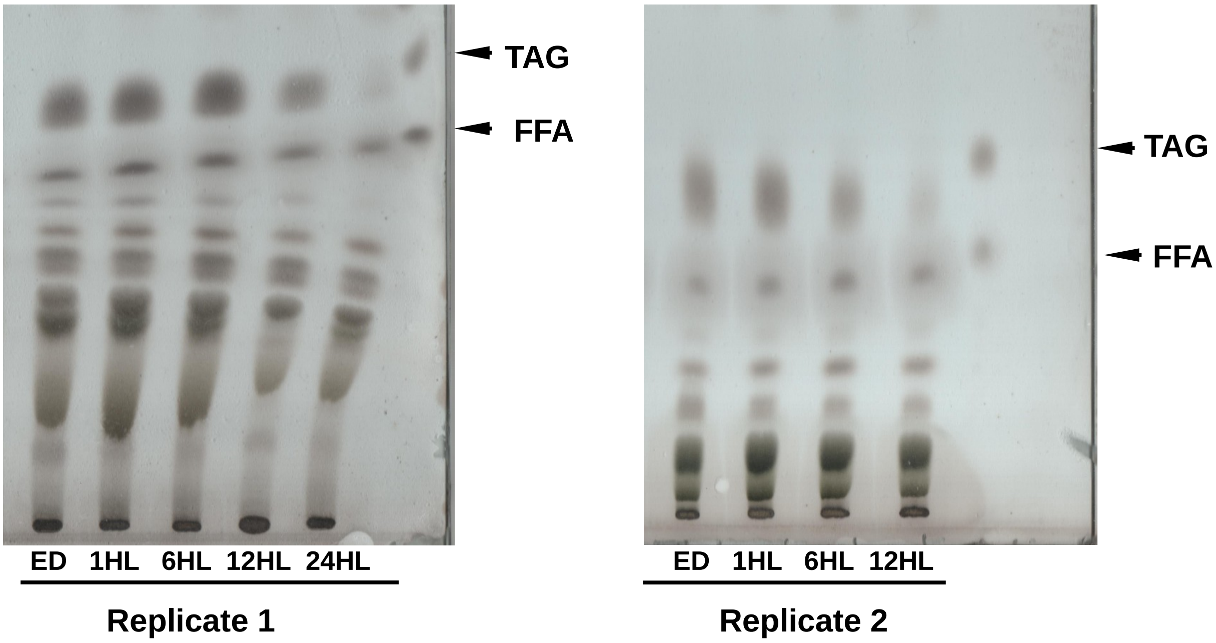


**E.** Strain cc-125, TAP media, 48 h of darkness (HD), then shifted to 15,000 lux of light (HL). Sample volumes corresponding to 4 million cells per lane were loaded.

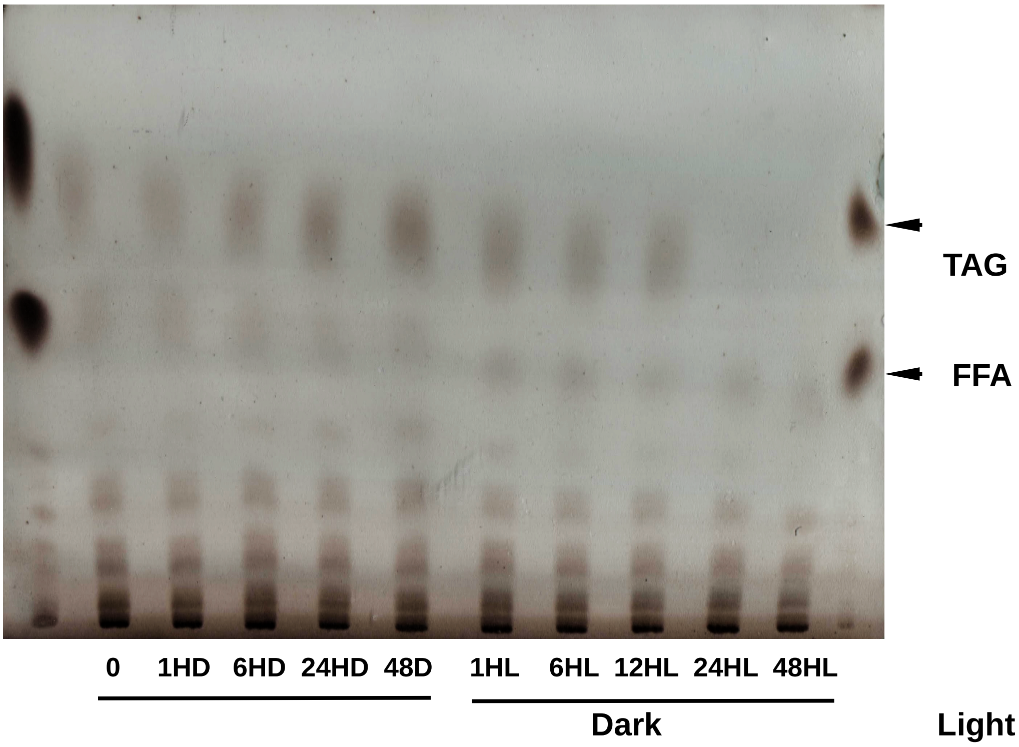

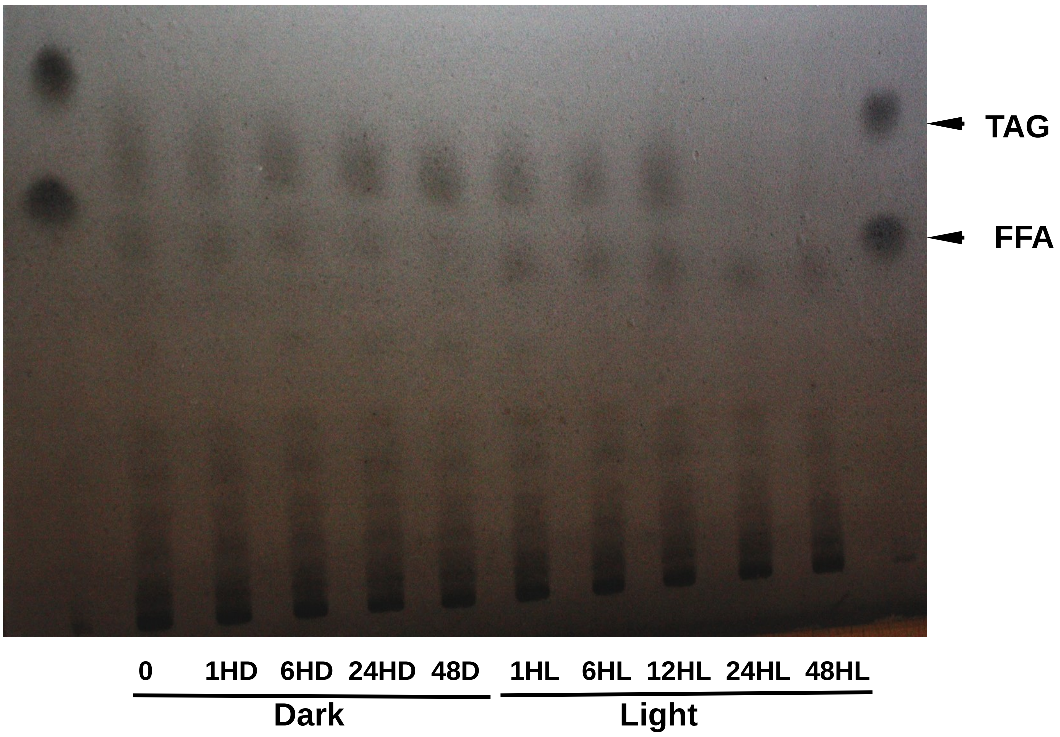


**F.** Strain cc-125, TAP media, 1,500 lux to 15,000 lux of light. Sample volumes corresponding to 10 million cells per lane were loaded.

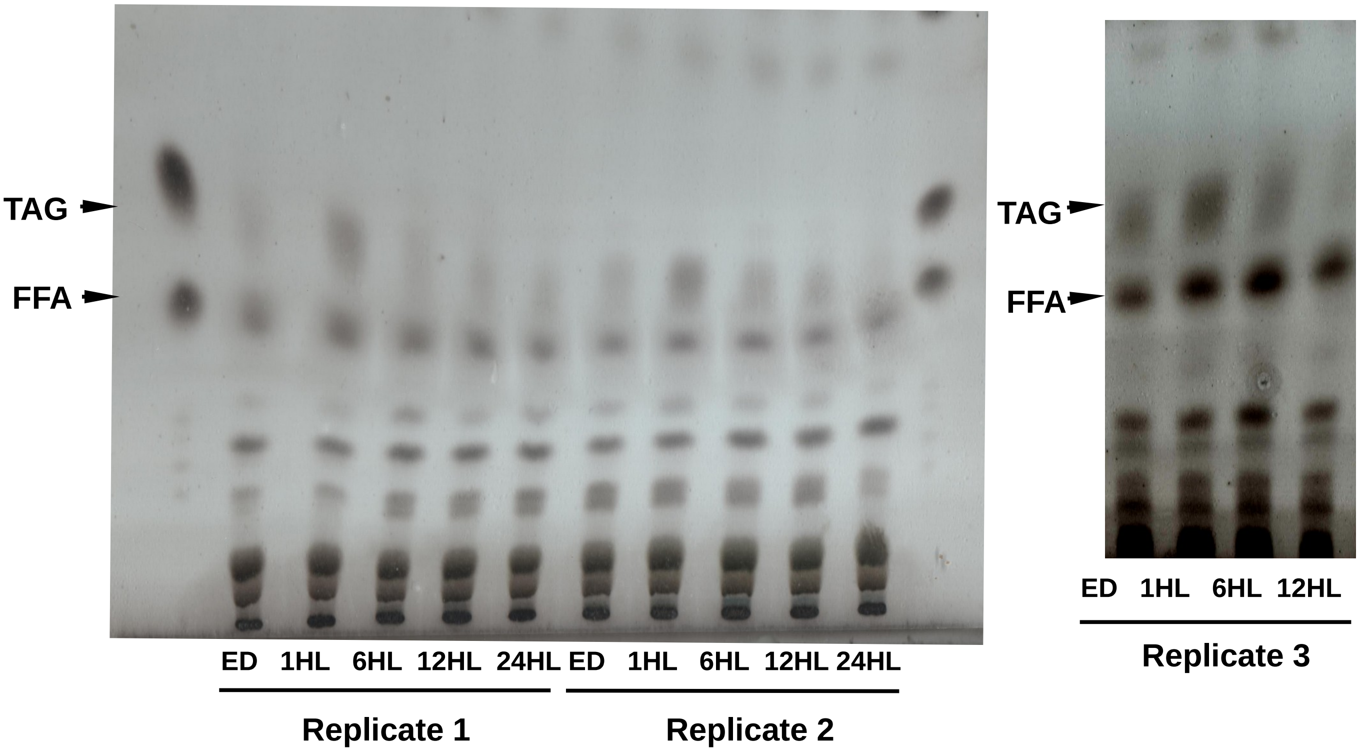
